# FMR1 KH0-KH1 Domains Coordinate m6A Binding and Phase Separation in Fragile X Syndrome

**DOI:** 10.1101/2025.03.17.643701

**Authors:** Xian Zhou, Chen-Jun Guo, Rui Wang, Yi-Lan Li, Tianyi Zhang, Zhuangyi Qiu, Shaorong Gao, Ji-Long Liu, Yawei Gao

**Author notes:** Correspondence to (SG); or (JLL); (YG).

## Abstract

The Fragile X Mental Retardation Protein (FMR1) regulates neurodevelopment via m6A RNA interactions, but domain-specific roles of KH0 and KH1 in RNA binding and pathology remain unclear. Through mutagenesis and AlphaFold3 modeling, we demonstrate that KH1 serves as the primary m6A-binding interface, whereas the KH0 domain (e.g., Arg138) modulates liquid-liquid phase separation (LLPS). Pathogenic mutations in KH0 disrupt RNA binding and enhance LLPS aggregation, while m6A RNA suppresses LLPS at KH0. Structural simulations reveal synergistic cooperation between KH0 and KH1 through hydrophobic and electrostatic networks. These dual-domain interactions establish a mechanistic link between m6A dysregulation, aberrant phase separation, and Fragile X syndrome pathogenesis. Our findings propose KH0 as a therapeutic target for RNA-driven neurodevelopmental disorders.

## INTRODUCTION

Fragile X syndrome, the most common form of inherited intellectual disability in humans, is an X-linked neurological disease that can be caused by genetic mutations or epigenetic modifications, resulting in the loss or reduction of its product, fragile X mental retardation protein (FMR1)^[1, 2]^. In addition, loss of the interaction between FMRP and RNA molecules can also lead to fragile X syndrome-related phenotypes^[3, 4]^.

As an RNA-binding protein, the amino acid sequence of FMR1 protein can be divided into three parts: N-terminal, KH domain, and C-terminal (containing RGG box) ^[5, 6]^. In addition, the presence of the KH0 region in front of the KH1 domain was recently discovered^[7, 8]^. Darnell et al had earlier identified 71 possible FMRP binding site sequences through the database. By constructing a series of FMR1 mutants, they found that the RGG box determines the binding of specific mRNA to the FMR1 protein. The KH domain serves as a regulatory motif that balances functional diversity and specificity, and therefore is widely used in biology^[9, 10]^.

An additional study showed that the FMR1 protein contains two KH-type and one RGG-type RNA-binding domains^[11, 12]^. I304 in the KH2 region is an important RNA-binding site. A patient with mental retardation carries a missense mutation (I304N). The idea that I304N may disrupt sequence-specific RNA recognition is also supported. Therefore, the mutual binding of KH2 domain and RNA was also confirmed.

The newly discovered KH0 region is located in the FMR1 amino acid sequence (126-202), and a missense mutation of R138 in this region was reported in a patient with ID and epileptic seizures. It shows that the KH0 region also plays an important role in the pathophysiology of FXS^[8]^. Although the domains of FMR1 KH0, KH1 and KH2 have been extensively studied, how they selectively bind and regulate the protein’s RNA targets remains unclear^[13, 14]^, and some of their functions in vitro also need to be further explored.

At the molecular level, m6A affects many aspects including the correct folding of mRNA, metabolism, translation, splicing, stability^[15]^ and the maturation of miRNA ^[16–18]^. The impact of m6A on RNA is mediated by an expanding array of m6A readers and m6A writers and potential erasers^[17, 19]^. Among them, the most studied m6A reader is the YTH domain family. YTHDF2 may select m6ARNA through the YTH domain and use the N-terminal domain to locate it on the P body, thus promoting the decay of mRNA^[20–22]^.

There is a recent report that FMR1 is another reader of m6A and reveals the link between mRNA modification and autism spectrum disorder^[22]^. As another reader of m6A, the ability of FMR1 protein to bind to m6ARNA, as well as the impact of a series of functions resulting from the interaction, have attracted widespread attention. According to reports in the literature, scientists used *Drosophila* as a model and explained that this combination causes dynamic phase switching of FMR1 particles, which ultimately leads to the decay of maternal mRNA and controls RNA processing and normal development of the body^[22, 23]^.

In this study, the mutant protein was obtained in vitro, and the changes in the binding force between the protein and RNA were determined through electrophoretic migration assay (EMSA) and surface plasmon resonance experiment (SPR). Microscopic imaging was used to observe the agglutination of protein particles, and electron microscopy was used to observe the particles in different Characteristics of the state.

We elucidate the role of key sites located in the core region of KH0 in RNA binding and phase separation, revealing a new important molecular site and domain for targeted intervention in FXS and autism. In addition, through experimental confirmation, ChimeraX software analysis and Alphafold3 prediction^[8, 24, 25]^, we proposed that methylation modification may change the folding method of RNA, leading to changes in its ability to bind to proteins, thereby regulating gene expression.

## RESULTS

### The KH0 domain interacts with nucleic acids

To understand the interaction between FMR1 protein and RNA, we obtained FMR1 protein and its mutant protein^[26]^ (Fs1) through gene recombination in vitro. We designed a pair of RNA probes (42 nucleotides in length)^[24, 27]^ and synthesized m6A modified sequences of the RNA probes. We conducted EMSA and SPR experiments by labeling probes with FAM and biotin. We obtained the following mutations: FMR1^F49N^, FMR1^R138Q^, FMR1^I304N^, FMR1^V308K^, FMR1^R534L&R546L^. These mutations encompass the important structural domain (Fs2) of FMR1 protein.

Using the FMR1^WT^ protein and these mutations, EMSA screening was performed to identify residues in the KH0 region where FMR1 binds to RNA and m6A modified RNA (**Figure 1A**). The FMR1^WT^ protein has strong interaction ability with RNA (ss-A) and m6A modified RNA (ss-m6A), while the mutant proteins FMR1^R534L&R546L^ and FMR1^F49N^ still have strong interaction ability with ss-A and ss-m6A. However, mutations in FMR1^R138Q^, FMR1^I304N^, and FMR1^V308K^ significantly affect the interaction with ss-m6A probes, but do not affect the interaction between FMR1 and ss-A probes.

**Figure 1.**
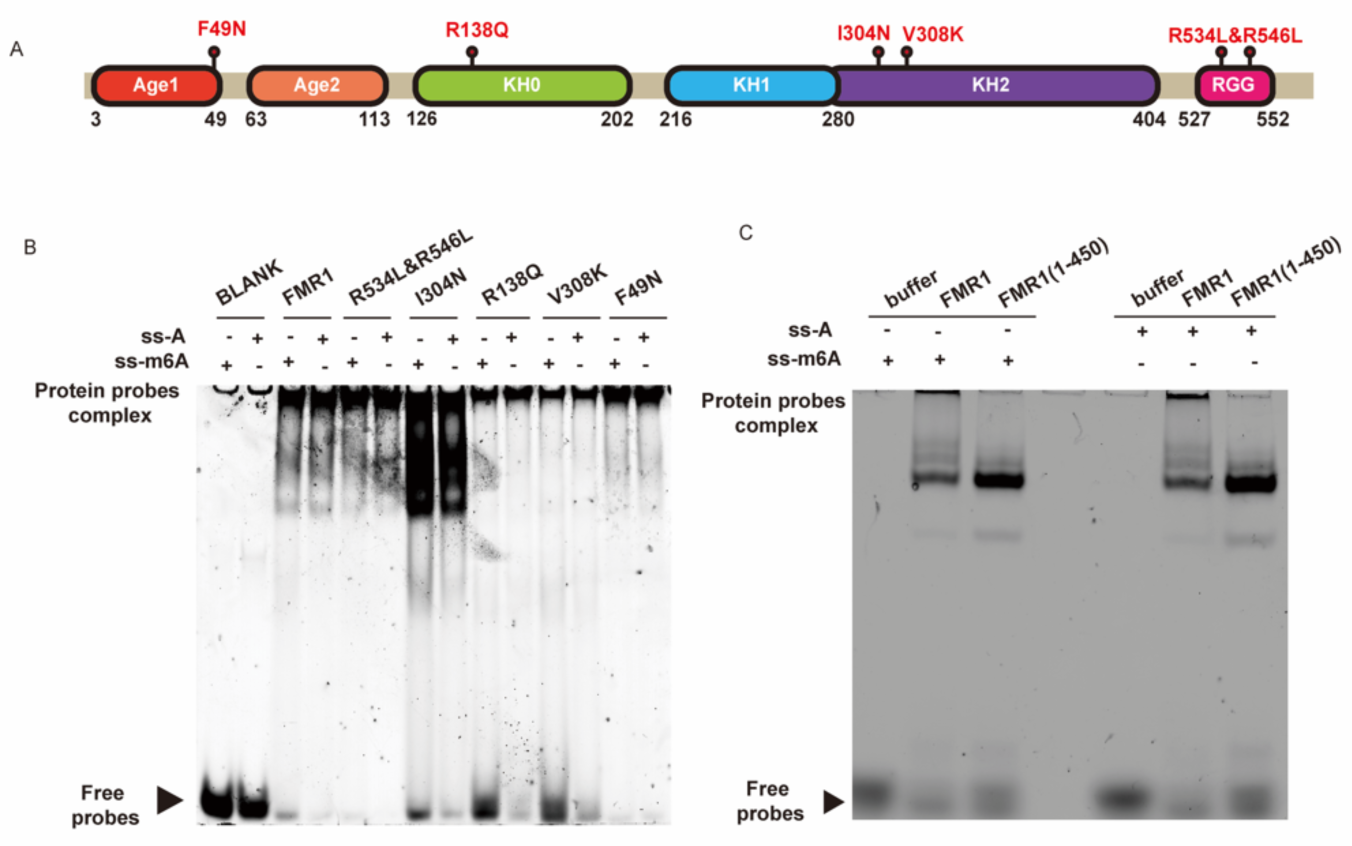
Detect the binding ability of wild-type or mutant FMR1 to m6A modified or unmodified RNA using EMSA. (A) Human FMR1 consists of six domains: Age1 (red), Age2 (orange), KH0 (green), KH1 (blue), KH2 (purple), and RGG box (rose red). The location of the mutations and domain boundaries are shown. (B) Perform EMSA to detect the binding affinity between wild-type or mutant FMR1 protein and RNA probes with or without m6A modification. (C) Perform EMSA to detect the binding affinity of full-length (FMR1) or truncated (FMR1 (1-450)) RNA probes with or without m6A modification.

Combining the EMSA experiments of FMR1^WT^ protein and FMR1(1-450) protein shown in **Figure 1B and C**, it can be found that the binding sites of RNA are distributed throughout the entire length of FMR1, or may have synergistic effects on different regions of the entire length when interacting with nucleic acid. FMR1^R138Q^, FMR1^I304N^, and FMR1^V308K^ are the newly identified key binding sites that interact with ss-m6A. Further confirmation was conducted using SPR experiments on the FMR1^WT^ protein and the R138 site located in the KH0 region. It was found that the binding ability of FMR1^WT^ to ss-m6A was stronger than that to ss-A (**Figure 2A and B**), and the R138Q mutation did indeed affect the interaction between FMR1 and ss-m6A probe (**Figure 2C and D**). Through microscopic observation, whether mutated or not, there is co localization between FMR1^R138Q^ and wild-type FMR1 and ss-A (**Figure 3**).

**Figure 2.**
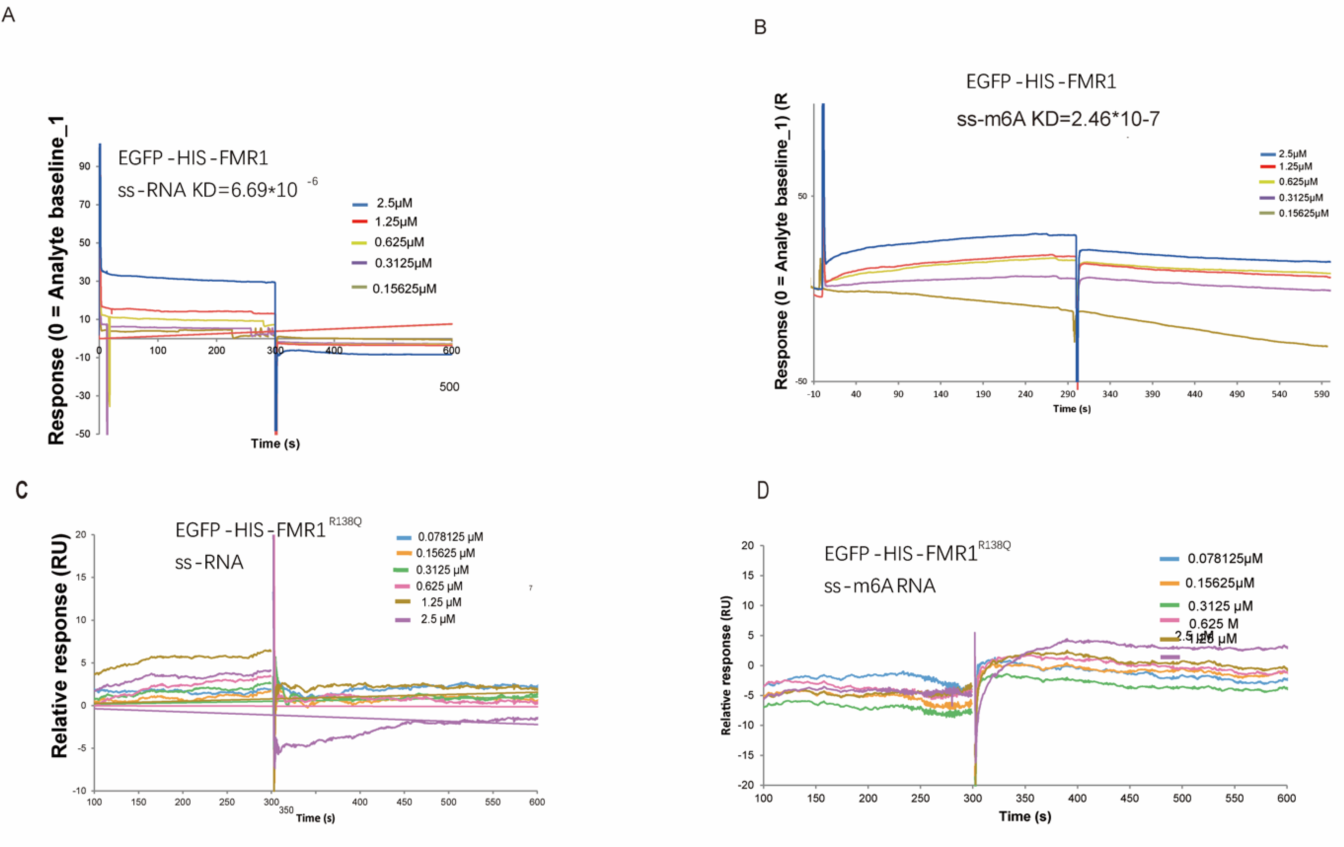
Detect the binding ability of wild-type or mutant FMR1 to m6A modified or unmodified RNA through SPR. (A, B) SPR assay is used to detect the binding affinity of FMR1WT protein to RNA probes, unmodified probes (A), or m6A modified probes (B). (C, D) SPR assay is used to detect the binding affinity of FMR1R138Q protein to RNA probes, unmodified RNA probes (C), or m6A modified probes (D). The equilibrium constant and kinetic constant are calculated by global fitting of the 1:1 Langmuir coupling model. The smaller the affinity (KD) value, the higher the affinity.

**Figure 3.**
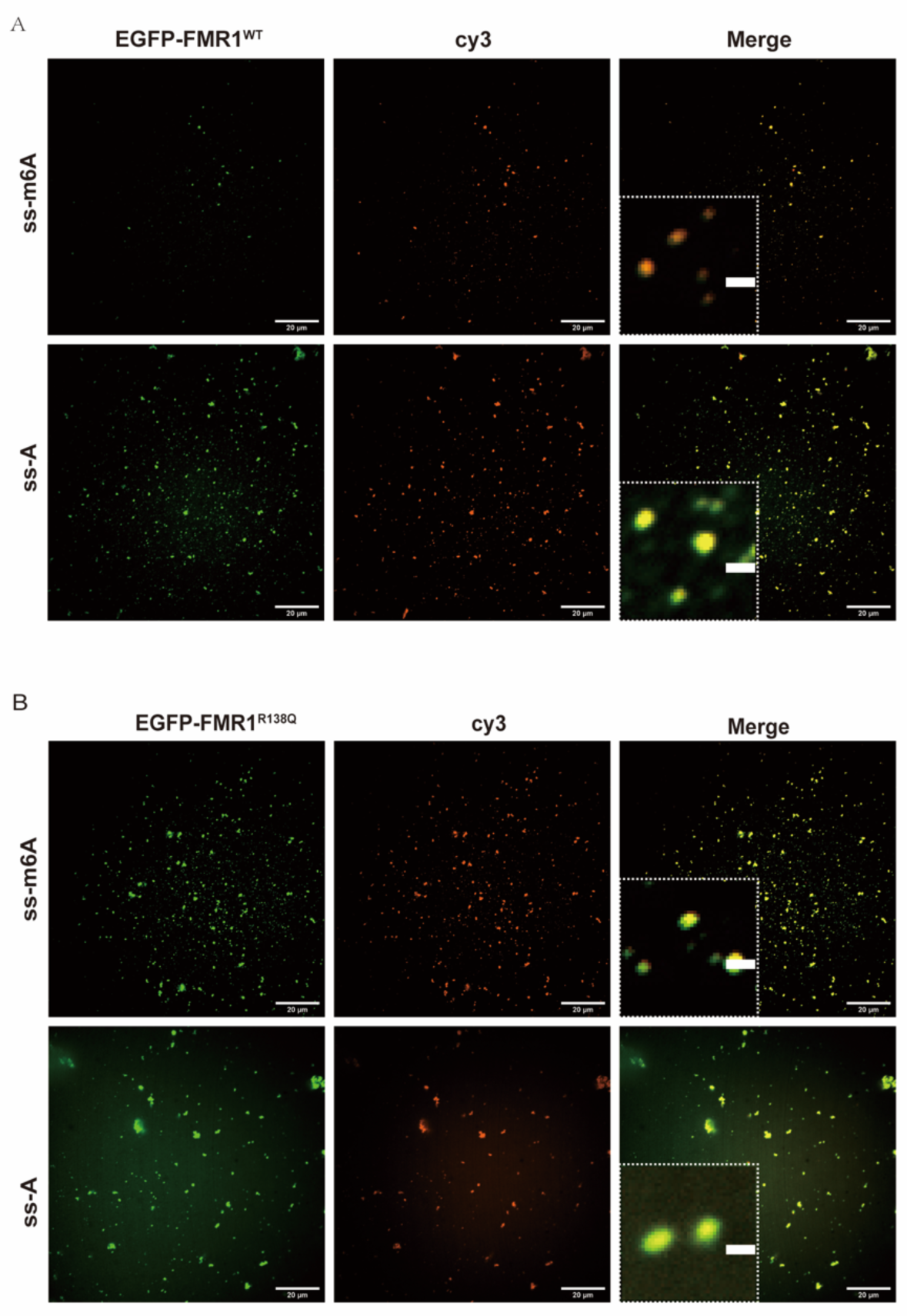
The location of the FMR1^WT^ or mutant FMR1^R138Q^ with m6A modified or unmodified RNA. (A, B) Both FMR1^WT^ and FMR1^R138Q^ proteins can form droplets, and contain m6A modified and unmodified RNAs in vitro.

These findings indicate that there is an interaction between ss-A and ss-m6A in the KH0 region of FMR1 protein, and there may be a direct site of action or a region that synergistically interacts with other sites. FMR1 preferentially binds to m6A modified RNA probes, and not only does the KH2 region have binding sites for m6A modified RNA, but site mutations in the KH0 region may also synergistically block the binding of FMR1 to m6A modified RNA. However, the R138Q mutation in the KH0 region does not affect the colocalization between the protein and nucleic acids.

### Interaction between FMR1 and RNA

In recent years, breakthroughs have been made in the field of protein prediction, with Alphafold3 being an AI model used for predicting proteins, RNA and DNA. Excellent effects have been demonstrated in the structure and interactions of biomolecules such as small molecules. We conducted structural comparative analysis and predicted the structure of RNA fragments and full-length FMR1 protein using Alphafold3 to explore the mode of action and binding site of fragile X intellectual disability protein (FMR1) with RNA.

The KH domain is a common motif for nucleic acid recognition in cellular functional proteins. The KH domain binds to RNA or ss RNA and is present in proteins related to transcription and translation regulation, as well as other cellular processes. We first compared the KH0 structure (4QVZ) of FMR1 with the RNA containing structures of three other KH family proteins (PDB Code: 1EC6, 5ELT, 2PY9)^[8, 10, 28]^, and the results showed that although KH0 has the typical β - α - α - β - β - α folding of the KH family. However, the α - helix1 and 2 positions of KH0 are offset relative to classical KH and are located in the RNA binding pocket of classical KH, which may limit the binding of RNA to it, suggesting that KH0 does not have the ability to bind to RNA (**Figure 4A-D**).

**Figure 4.**
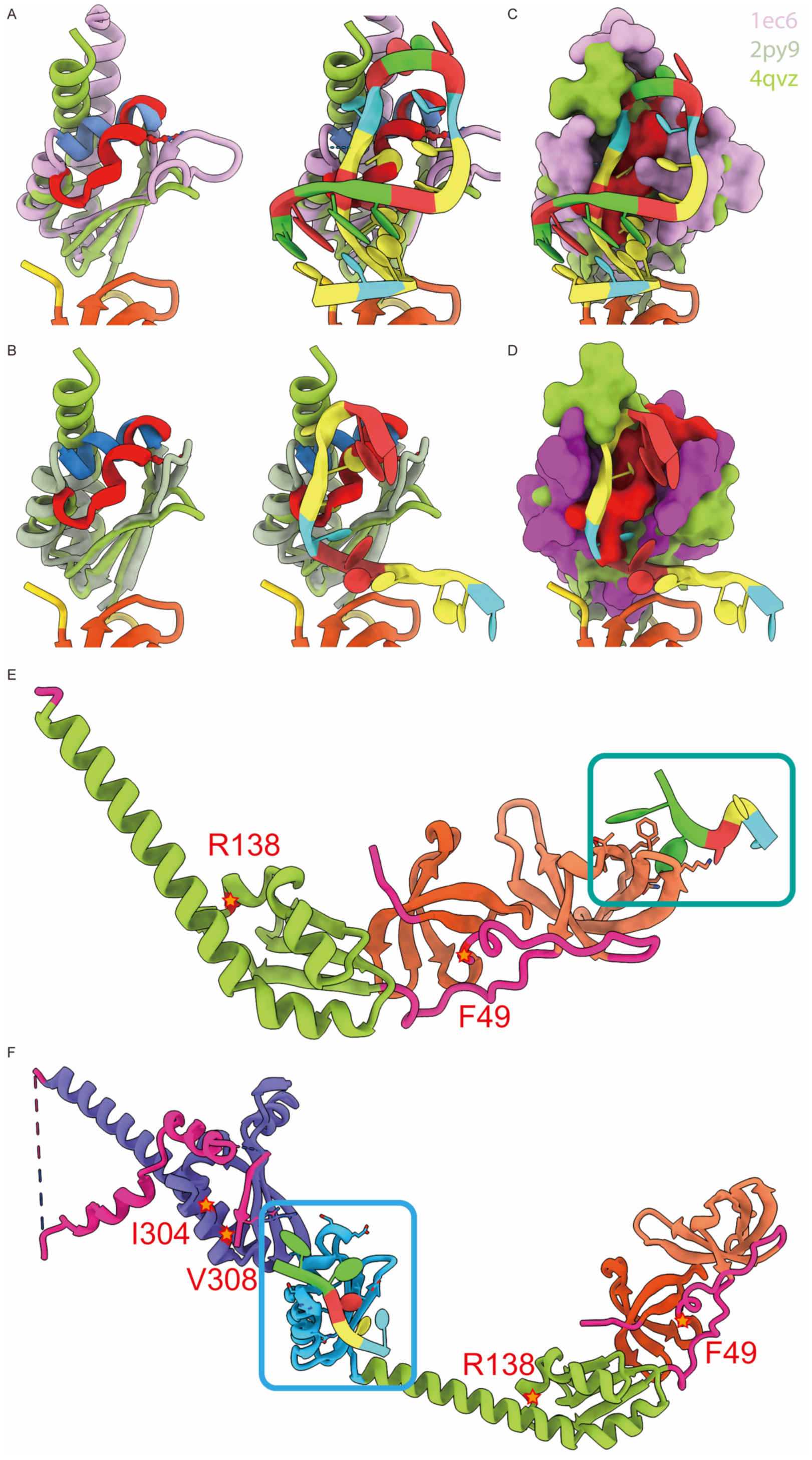
The structural comparison and Alphafold3 prediction display of FMR1 proteins. (A) The comparison between the 4QVZ of FMR1 containing the KH domain (green) and the KH family protein 1EC6 (pink) reveals the alpha helix in both domains. (B) Comparison of 4QVZ and 1EC6 structures with RNA, as well as the electrostatic surface of the KH domain and RNA complex. The KH0 domain is located in the binding pocket between KH and RNA. (C) The comparison between the 4QVZ of FMR1 containing the KH domain (green) and the KH family protein 2PY9 (gray green) reveals the alpha helix in both domains. (D) Comparison of 4QVZ and 2PY9 structures with RNA, as well as the electrostatic surface of the KH domain and RNA complex. The KH0 domain is located in the binding pocket between KH and RNA. (E) The position map of the amino acid domains (1-215) of RNA simulated by FMR1 and Alphafold3, with blue boxes indicating possible contact positions. (F) The location map of full-length FMR1 and RNA simulated by Alphafold3. The blue box indicates possible contact locations.

Subsequently, we used Alphafold3 to predict the eutectic structure between the nucleic acid sequence (1-215) of FMR1 and the key RNA: GGACU, and observed the spatial position between the protein and RNA to speculate whether they have a basis for interaction. The predicted FMR1 (1-215) as shown in **Figure 4E** has a similar morphology to 4QWZ. But RNA binds to the end of Age2 and not to KH0.

In addition, using the full-length sequence of FMR1 and nucleic acid for complex structure prediction, the results showed (**Figure 4F**) that RNA binds to the KH1 domain. The above analysis and simulation experiment results indicate that the KH0 region does not seem to have the conditions for direct interaction with RNA.

In vitro mutations FMR1^F49N^ (Age1 region), FMR1^R138Q^ (KH0 region), FMR1^I304N^ (KH2 region), FMR1^V308K^ (KH2 region), and FMR1^R534L&R546L^ (RGG box) are not located in the KH1 region, so they may not have affected the interaction between the mutant protein and RNA. However, due to the methylation of RNA, in vitro experiments have found that methylated RNA has a reduced binding ability to FMR1^R138Q^, FMR1^I304N^, and FMR1^V308K^ mutant proteins. This indicates that methylated RNA has an impact on the KH0 and KH2 regions.

Since Alphafold3 cannot simulate the interaction between methylated RNA and proteins, we speculate that the reasons affecting the binding ability between nucleic acid and FMR1 protein may not only be due to direct binding sites, but also because methylation changes the folding mode of RNA, or may change the synergistic effect between different structural domains. The two mutation sites located in the KH2 region, FMR1^I304N^ and FMR1^V308K^, may be due to a change in surface charge, leading to an imbalance of charges in the hydrophobic network and disrupting the protein conformation.

### Phase separation of FMR1

Abnormal phase separation in living organisms may lead to the occurrence of diseases. In the droplets formed by phase separation, some proteins can condense to form harmful solid substances, causing diseases. Disease treatment can be achieved through interference, control, and disruption of phase separation. Protein phase separation is of great significance for cellular tissue and signal transduction^[29–31]^. The FMR1 protein is the main component of dynamic neuronal granules transported from somatic cells to dendrites^[32–34]^, and it belongs to RNA binding proteins. Its main features include a hydrophobic network, an interface for pi-pi interactions, and a high content of arginine, which is predicted to have phase separation related characteristics^[35]^.

We attempted to observe the changes in protein particle size and differences in polymerization ability after co localization between proteins and nucleic acids in vitro, in order to observe whether proteins form phase separation. Through confocal microscopy, it was found that the FMR1^WT^ protein undergoes coagulation when it reaches a certain concentration in solution. After adding ss-A and ss-m6A, ss-A significantly increased the size of FMR1^WT^ protein RNA droplets. In contrast, the m6A modified RNA probe did not significantly alter the droplet size of FMR1 (**Figure 5**). Fluorescence recovery analysis after photobleaching showed that compared with droplets formed by FMR1^WT^ protein with ss-A and ss-m6A and droplets containing only FMR1^WT^ protein, droplets of FMR1^WT^ and ss-m6A had the least recovery after photobleaching (**Figure 6A and B**), indicating that protein binding with ss-m6A significantly altered the fluidity of FMR1^WT^ protein condensate.

**Figure 5.**
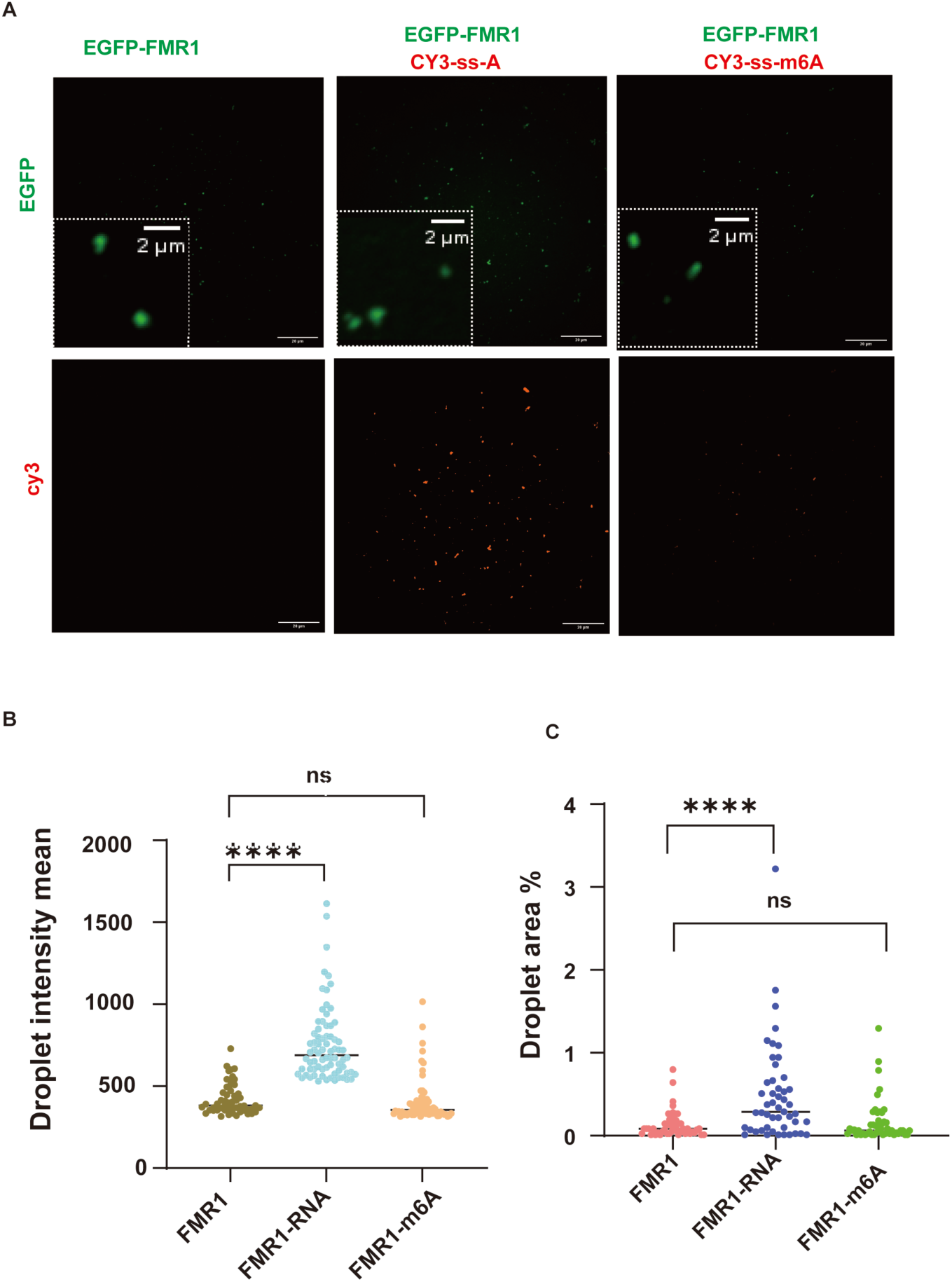
Formation of FMR1^WT^ protein droplets after adding labeled RNA in vitro. (A) Droplet formation analysis of GFP-FMR1^WT^ mixed with unmodified probe or Cy3 labeled m6A modified probe. Scale bar, 5 μ m. (B) Quantitative area of green droplet signal in (a) (GFP-FMR1^WT^+Cy3-ss-a, P<0.0001; GFP-FMR1^WT^+Cy3-s-m6A, P=0.47). (C) Quantitative intensity of green droplet signal in (a) (GFP-FMR1^WT^+Cy3-ss-A, (* * * * P<0.0001; GFP-FMR1^WT^+Cy3-s-m6A, P=0.37).

**Figure 6.**
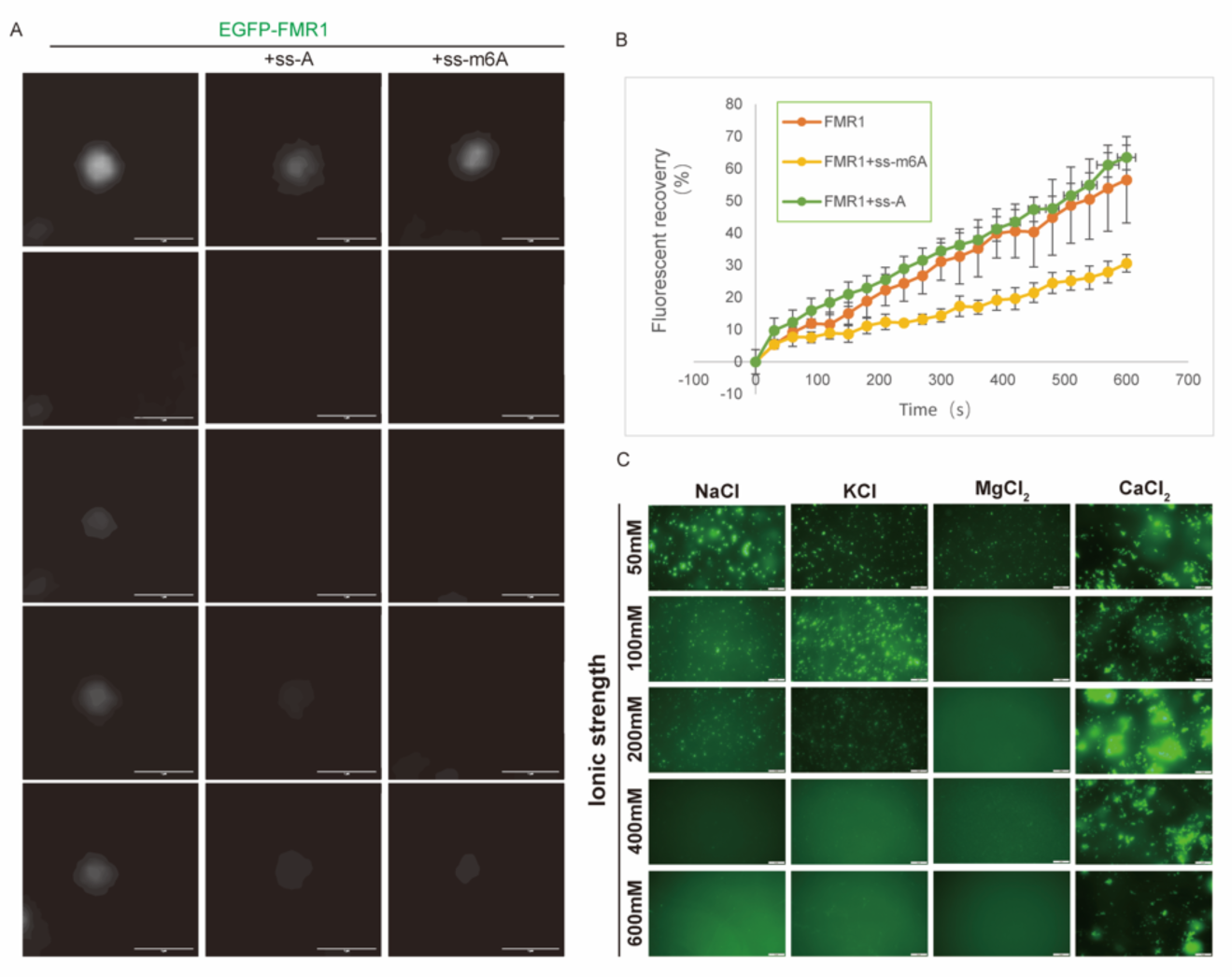
m6A modification on FMR1^WT^ fluidity and the effect of salt ion concentrations on droplet formation. (A) Fluorescence recovery analysis after photobleaching to study the flowability of FMR1^WT^ droplets. Three independent images were displayed, along with images of droplet photobleaching at different time points. Scale bar, 1 μ m. (B) We plotted the fluorescence intensity of unmodified probes or Cy3 labeled m6A modified RNA droplets over time after photobleaching. This curve represents the average percentage of fluorescence recovered after photobleaching of different droplets (n=3). The error bar represents the mean ± standard deviation. (C) FMR1^WT^ images at the same concentration of 10 μ M protein, 25 μ M Na2PO4 pH 7.4, 2 mM DTT, and different ion concentrations.

We previously confirmed that FMR1 phase separation does not require any additives, and we attempted to interfere with the electrostatic interactions present within FMR1^WT^ protein droplets. By observing phase diagrams under different protein and salt concentrations using a microscope, we found that an increase in salt concentration is not conducive to the formation of FMR1 ^WT^ protein droplets^[36]^.

To test whether this result is caused by general ion strength effects or specific salt ion effects, we incubated FMR1^WT^ protein droplets in salt solutions of different valences (KCl, MgCl2, CaCl2, NaCl) and found that the degree of damage to droplet formation varied among different salt solutions (**Figure 6C**), with divalent ions more likely to disrupt droplet formation. Therefore, ion strength can regulate the formation ability of FMR1^WT^ protein droplets in vitro, which may be the result of electrostatic interactions. It is worth noting that the addition of Ca^2+^ changes the morphology of protein aggregation, indicating that Ca^2+^ may affect the activity or internal structure of proteins. It is also possible that Ca^2+^ strongly positively affects the electrical interactions between some amino acids and nucleic acids.

At the same time, we found that the electrostatic force of this ion still exists in the presence of nucleic acid (Fs3). Our work shows that FMR1 protein functions in an oligomeric state, and m6A modified RNA inhibits the formation of higher-order protein assembly, allowing translation to proceed normally. Salt ions can alter the protein’s ability to form phase separation, and the higher the salt ion concentration, the weaker the phase separation phenomenon.

### *FMR1^R138Q^* reduces the interaction with ss-m6A

An m6A RNA has the ability to bind to FMR1 protein and affects the droplet fluidity of protein formation phase separation. There are reports of missense mutations in the FMR1 protein from arginine 138 to glutamine (R138Q) in a patient with intellectual disability and epileptic seizures^[37, 38]^. The R138Q site is located in the KH0 region, which has little research on its properties and functions in the body. The function of FMR1^R138Q^, as a key site in diseases, is worth exploring. Through confocal microscopy, it was found that the FMR1^R138Q^ mutant protein and FMR1^WT^ can form droplets on their own at the same concentration in 50mM NaCl, and the FMR1^R138Q^ mutant protein can significantly enhance the protein phase separation ability, resulting in a significant increase in droplet size and fluorescence intensity of the mutant protein in solution (**Figure 7A-C**). After adding ss-A and ss-m6A, there was no significant change in droplet size, but there was a significant change in fluorescence intensity. Among them, ss-m6A significantly enhanced the fluorescence intensity of the mutant protein FMR1^R138Q^ (**Figure 7D-F**).

**Figure 7.**
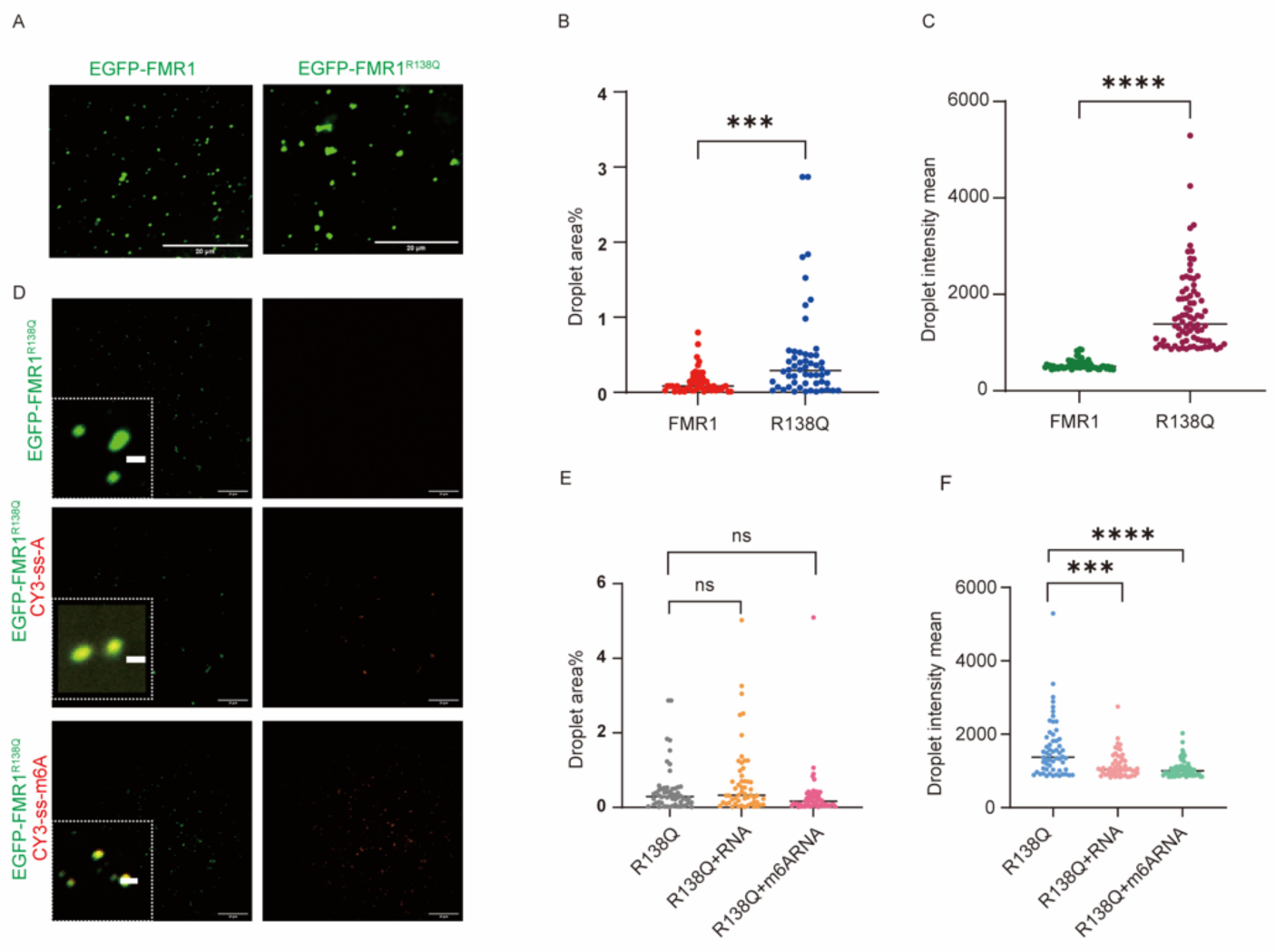
The effect of FMR1^R138Q^ on droplet size. (A) Comparison of droplets of FMR1^R138Q^ and GFP-FMR1^WT^ without unmodified RNA and m6A RNA. The scale bar is 5 μ m. Quantitative area and intensity of droplet signals in (B, C) (a) (* * * P<0.0001, * * * * P<0.0001). (D) Droplet formation assay of GFP-FMR1^R138Q^ mixed with unmodified RNA probe or Cy3 labeled m6A modified RNA probe. Scale bar, 5 μ m. (E) Quantitative area of green droplet signal in (d) (GFP-FMR1^R138Q^ + Cy3-ss-A, P =0.13; GFP-FMR1^R138Q^ + Cy3-ss-m6A, P = 0.17). (F) Quantitative intensity of green droplet signal in (d) (GFP-FMR1^R138Q^ + Cy3-ss-A, (***P <0.0002; GFP-FMR1^R138Q^ + Cy3-ss-m6A, ****P <0.0001).

Fluorescence recovery analysis after photobleaching showed that the FMR1^R138Q^ mutation significantly increased the fluidity of the protein (**Figure 8**), while the addition of ss-A and ss-m6A rapidly reduced the fluidity of the mutated protein FMR1^R138Q^. In summary, FMR1^R138Q^ reduced the interaction between the protein and ss-m6A, and this change in binding ability led to more pronounced droplet aggregation of the protein. The addition of ss-A and ss-m6A can cause protein depolymerization.

**Figure 8.**
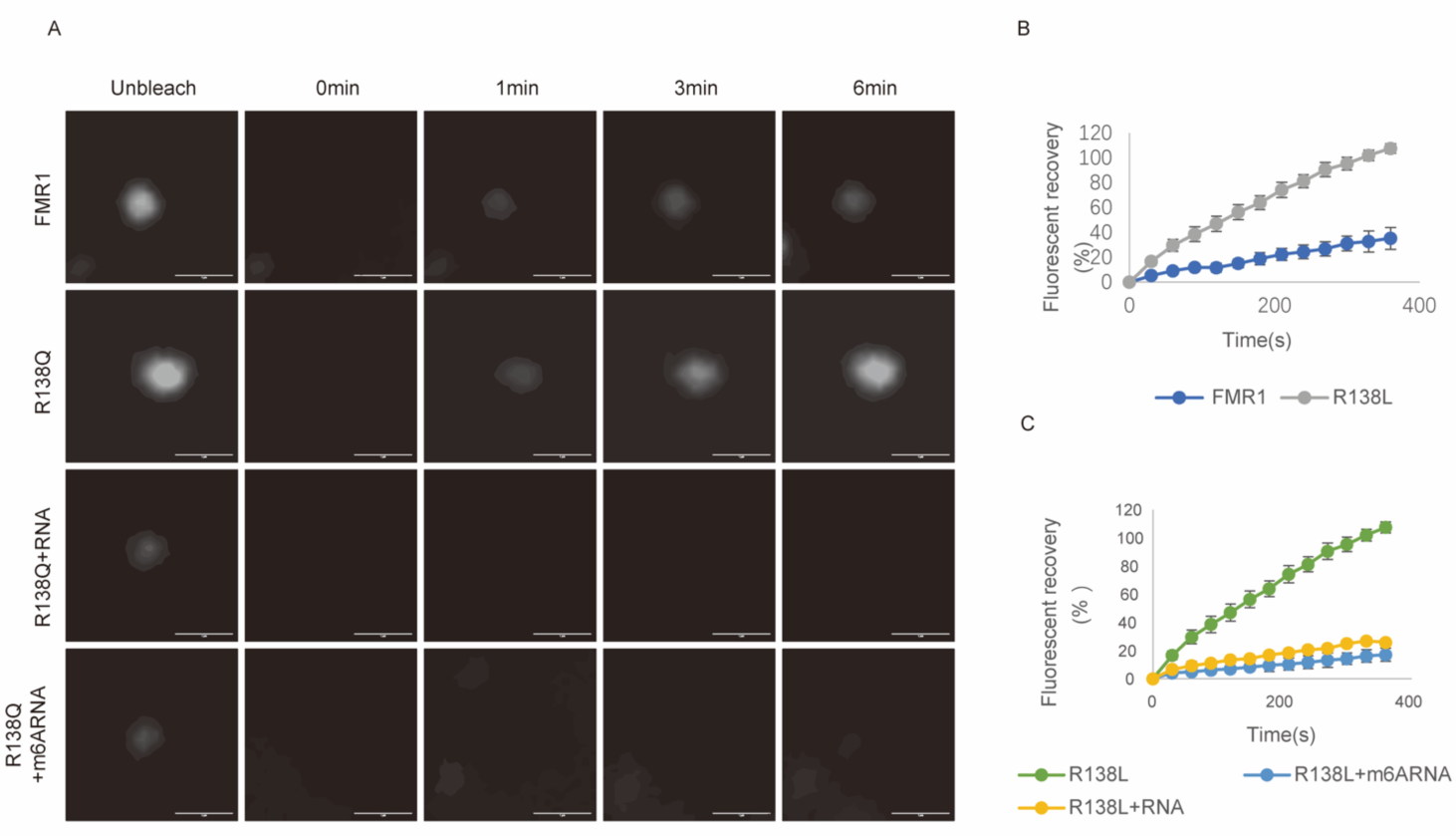
The effect of FMR1^R138Q^ protein phase separation. (A) Fluorescence recovery after photobleaching assays to study the fluidity of GFP-FMR1^WT^ and GFP-FMR1^R138Q^ droplets without RNA, and GFP-FMR1^R138Q^ droplets add unmodified RNA or m6A RNA. Display images of droplet photobleaching at different time points. Scale bars, 1μm. (B) Recovery of GFP-FMR1WT and GFP-FMR1R138Q droplets fluorescence intensity over time after photobleaching was plotted. The curve represents the average percentage of fluorescence recovered after photobleaching of different droplets (n=3). Error bars represent mean ± SD. (C) Recovery of GFP-FMR1R138Q droplet fluorescence intensity with unmodified RNA or m6A RNA over time after photobleaching was plotted. The curve represents the average percentage of fluorescence recovered after photobleaching of different droplets (n=3). The error bars represent mean ± SD.

## DISCUSSION

Our study shows that in the full-length sequence of FMR1^WT^, there is a motif for preferential binding of methylation-modified RNA. Mutations in these key sites also have disease manifestations in some patients with Fragile X Syndrome (FXS), indicating that sites that preferentially bind methylation-modified RNA are more closely related to the disease. At the same time, we also found that human FMR1^WT^ itself has phase separation phenomenon under a certain salt concentration.

The experiment uses the *E. coli* system to express full-length human EGFP-HIS-FMR1, and purifies the target protein through affinity chromatography and molecular sieves. The purified EGFP-HIS-FMR1 protein has complete biochemical activity, which is particularly important for in vitro protein research. step. In the case of complex in vivo environment, conducting experiments by simulating the in vivo environment in vitro is of guiding significance for initially exploring the important properties of proteins. FMR1 has been studied as an RNA-binding protein for many years. In recent studies, the connection between *Drosophila* FMR1 and methylation-modified RNA and its relationship with phase separation have been deeply explored ^[8]^. As human FMR1, in-depth research on it will better connect it with the treatment of related diseases.

Phase separation plays an important role in the cytoskeleton, gene activation, gene assembly, and transcription. Abnormal phase separation can lead to diseases. For example, the proteins in the droplets formed by phase separation will condense and harden, forming harmful solid substances, leading to the occurrence of diseases. Disease treatment can be achieved by interfering, controlling, and destroying phase separation. We use the newly discovered pathogenic site arginine 138 mutation in the KH0 region as the research object and obtained humanFMR1^R138Q^ in vitro. We find that the protein shows stronger phase separation than FMR1^WT^ in vitro, and find that RNA interferes with the phase separation of the protein. Further, we find that salt ion strength is a key factor in the formation of phase separation. Calcium ions have a greater impact on the protein structure, causing a large amount of protein to precipitate.

The KH motif consists of approximately 70 amino acids, and KH domains are identified as nucleic acid recognition motifs in proteins that perform a wide range of cellular functions. We are surprised to find that the in vitro biochemical experiment results are inconsistent with the prediction results of Alphafold3 and the comparison results of the software. Alphafold3 shows that the protein region recognized by nucleic acids is mainly in the KH1 region. R138 is located on the α-helix of the KH0 region of the FMR1 protein and does not itself have the ability to bind to nucleic acids, the mutation from R138 to Q only changes the charge but does not change the folding mode of the KH0 region.

However, biochemical experiments show that the R138 site affects the interaction between m6ARNA and protein. We speculate that the interaction between FMR1 protein and nucleic acid may not only be directly affected by the site, but also caused by the synergistic interaction of multiple regions. In vitro biochemical experiments show that methylated RNA affects the ability of some sites on the FMR1 protein to interact with RNA, which may be the reason for the change in charge around the protein.

The living body is a complex environment, and the causes of Fragile beyond the site, there are still many factors that contribute to the occurrence of the disease. These in vitro experiments and prediction experiments provide new ideas for us to gradually understand the functions of FMR1 in the body and the impact of methylated nucleic acids on the function of the nucleic acid-binding protein FMR1.

## MATERIALS AND METHODS

### Expression and Purification of FMR1 and Mutant Proteins

The full-length human FMR1 gene was cloned into a vector with an N-terminal 6XHis-tag and an EGFP tag under the control of a lac1 promoter. The construct was transformed into Transetta (DE3) cells for protein expression. Induction was performed using 1 mM IPTG at 14°C for over 18 hours. Cells were harvested by centrifugation at 4000 rpm for 15 minutes and resuspended in cold lysis buffer (500 mM NaCl, 50 mM Tris-HCl [pH 8.0], 10% glycerol). After lysis, the supernatant was collected by centrifugation at 18,000 rpm for 60 minutes at 4°C. The supernatant was incubated with Ni-Agarose resin (Qiagen) for 1 hour, followed by washing with lysis buffer supplemented with 40 mM imidazole. The protein was eluted using lysis buffer containing 250 mM imidazole and 1 mM β-mercaptoethanol (β-Me) at pH 8.0. The purified FMR1 protein was concentrated to approximately 2 mg/mL and stored in a buffer containing 250 mM NaCl and 25 mM Tris-HCl (pH 8.0).

### Electrophoretic Mobility Shift Assays (EMSAs)

To identify RNA-binding residues and assess binding affinity, FAM-labeled single-stranded RNA probes (FAM-AUGGGCCGUUCAUCUGCUAAAAGGXCUGCUUUUGGGGCUUGU-3’, where X = A or m6A) were synthesized by Tsingke Biotechnology. Purified FMR1 protein was incubated with the RNA probes in a reaction buffer (25 mM Tris-HCl [pH 8.0], 150 mM NaCl, 50 mM KCl, 5 mM MgCl2, 10 mM DTT, 1 μM BSA) for 30 minutes at room temperature. The RNA-protein complexes were resolved on an 8% native polyacrylamide gel in 0.5× TBE buffer at 120 V for 35 minutes. Gels were visualized using a Typhoon FLA9500 scanner (GE Healthcare)^[39]^.

### Surface Plasmon Resonance (SPR) Measurements

Biotinylated RNA probes (biotin-UGGGCCGUUCAUCUGCUAAAAGGXCUGCUUUUGGGGCUUGU-3’, where X = A or m6A) were synthesized by Tsingke Biotechnology and immobilized on SA chips (Cytiva). Binding kinetics were analyzed using a Biacore 8K instrument (GE Healthcare). Increasing concentrations of FMR1 protein were injected over the chip surface at a flow rate of 30 µL/min for 120 seconds, followed by a 300-second dissociation phase in running buffer (1× PBS, 0.1% BSA, 0.02% Tween 20) at 25°C. Binding constants were calculated using the 1:1 Langmuir binding model with Biacore 8K evaluation software.

### Phase Separation Assay

GFP-tagged FMR1WT and FMR1R138Q proteins were expressed and purified from E. coli. Phase separation experiments were performed by mixing 10 µM protein with a buffer (20 mM Tris-HCl [pH 8.0], 50 mM NaCl, 25 mM Na2HPO4). For RNA-dependent phase separation, 10 µM protein was incubated with 5 µM Cy3-labeled A/m6A RNA probes in the same buffer. Samples were immediately transferred to a glass-bottom dish and imaged using a Nikon CSU SORA confocal microscope with a 60× oil immersion lens (488 nm and 405 nm lasers). Differential interference contrast (DIC) images were acquired using an OLYMPUS ix73 microscope with a 100× oil immersion lens.

### Fluorescence Recovery After Photobleaching (FRAP)

To assess droplet fluidity, FMR1 or mutant protein droplets were photobleached using a 488 nm laser (50% intensity) on a Nikon CSU SORA confocal microscope. Images were captured every 30 seconds for up to 6 minutes. Fluorescence recovery was quantified by comparing the intensity of the bleached area to an unbleached region of the same size. Recovery ratios were calculated from at least five spots across three independent samples.

## ACKNOWLEDGMENTS

We thank Zhi Zhou for guiding the phase separation experiments. We thank the Molecular and Cell Biology Core Facility and Core Imaging Facility at the School of Life Science and Technology. We also thank Shanghai Institute for Advanced Immunochemical Studies (SIAIS). This work was primarily supported by the National Key R&D Program of China (2021YFA0804700 to J.-L.L., 2024YFA1107000 to Y.G., and 2022YFC2702200 to S.G.), the National Natural Science Foundation of China (32370744 and 32350710195 to J.-L.L., 32370869 to Y.G., and 92168205 and 32330030 to S.G.), Medical Research Council (MC_UU_12021/3 and MC_U137788471 to J.-L.L.), the Shanghai Pilot Program for Basic Research (to Y.G.), and the Peak Disciplines (type IV) of Institutions of Higher Learning in Shanghai (to S.G.).

## REFERENCES

[1] Bagni C, Tassone F, Neri G, et al. Fragile X syndrome: causes, diagnosis, mechanisms, and therapeutics [J]. J Clin Invest, 2012, 122(12): 4314–22.

[2] Bagni C, Zukin R S. A Synaptic Perspective of Fragile X Syndrome and Autism Spectrum Disorders [J]. Neuron, 2019, 101(6): 1070–88.

[3] Ashley C T, Jr., Wilkinson K D, Reines D, et al. FMR1 protein: conserved RNP family domains and selective RNA binding [J]. Science, 1993, 262(5133): 563-6.

[4] Verkerk A J, Pieretti M, Sutcliffe J S, et al. Identification of a gene (Fmr-1) containing a CGG repeat coincident with a breakpoint cluster region exhibiting length variation in fragile X syndrome [J]. Cell, 1991, 65(5): 905–14.

[5] Siomi H, Siomi M C, Nussbaum R L, et al. The protein product of the fragile X gene, FMR1, has characteristics of an RNA-binding protein [J]. Cell, 1993, 74(2): 291–8.

[6] Brown V, Small K, Lakkis L, et al. Purified Recombinant FMRP exhibits selective RNA binding as an intrinsic property of the fragile X mental retardation protein [J]. J Biol Chem, 1998, 273(25): 15521–7.

[7] Adinolfi S, Bagni C, Musco G, et al. Dissecting FMR1, the protein responsible for fragile X syndrome, in its structural and functional domains [J]. RNA, 1999, 5(9): 1248–58.

[8] Myrick L K, Hashimoto H, Cheng X, et al. Human Fmrp contains an integral tandem Agenet (Tudor) and KH motif in the amino terminal domain [J]. Hum Mol Genet, 2015, 24(6): 1733–40.

[9] Nicastro G, Taylor I A, Ramos A. KH-Rna interactions: back in the groove [J]. Curr Opin Struct Biol, 2015, 30: 63–70.

[10] Valverde R, Edwards L, Regan L. Structure and function of KH domains [J]. Febs j, 2008, 275(11): 2712–26.

[11] Darnell J C, Fraser C E, Mostovetsky O, et al. Kissing complex RNAs mediate interaction between the Fragile-X mental retardation protein KH2 domain and brain polyribosomes [J]. Genes Dev, 2005, 19(8): 903–18.

[12] Vasilyev N, Polonskaia A, Darnell J C, et al. Crystal structure reveals specific recognition of a G-quadruplex RNA by a β-turn in the RGG motif of FMRP [J]. Proc Natl Acad Sci U S A, 2015, 112(39): E5391–400.

[13] Ascano M, Jr., Mukherjee N, Bandaru P, et al. FMRP targets distinct Mrna sequence elements to regulate protein expression [J]. Nature, 2012, 492(7429): 382–6.

[14] Greenblatt E J, Spradling A C. Fragile X mental retardation 1 gene enhances the translation of large autism-related proteins [J]. Science, 2018, 361(6403): 709–12.

[15] Wu B, Li L, Huang Y, et al. Readers, writers and erasers of N(6)-methylated adenosine modification [J]. Curr Opin Struct Biol, 2017, 47: 67–76.

[16] Meyer K D, Jaffrey S R. Rethinking m(6)A Readers, Writers, and Erasers [J]. Annu Rev Cell Dev Biol, 2017, 33: 319–42.

[17] Roundtree I A, Evans M E, Pan T, et al. Dynamic Rna Modifications in Gene Expression Regulation [J]. Cell, 2017, 169(7): 1187–200.

[18] Visvanathan A, Somasundaram K. mrna Traffic Control Reviewed: N6-Methyladenosine (m(6) A) Takes the Driver’s Seat [J]. Bioessays, 2018, 40(1).

[19] Zaccara S, Ries R J, Jaffrey S R. Reading, writing and erasing mrna methylation [J]. Nat Rev Mol Cell Biol, 2019, 20(10): 608–24.

[20] Lee M, Kim B, Kim V N. Emerging roles of Rna modification: m(6)A and U-tail [J]. Cell, 2014, 158(5): 980–7.

[21] Wang X, Lu Z, Gomez A, et al. N6-methyladenosine-dependent regulation of messenger Rna stability [J]. Nature, 2014, 505(7481): 117–20.

[22] Edupuganti R R, Geiger S, Lindeboom R G H, et al. N(6)-methyladenosine (m(6)A) recruits and repels proteins to regulate mrna homeostasis [J]. Nat Struct Mol Biol, 2017, 24(10): 870–8.

[23] Zhang G, Xu Y, Wang X, et al. Dynamic FMR1 granule phase switch instructed by m6a modification contributes to maternal Rna decay [J]. Nat Commun, 2022, 13(1): 859.

[24] Liao S, Sun H, Xu C. Yth Domain: A Family of N(6)-methyladenosine (m(6)A) Readers [J]. Genomics Proteomics Bioinformatics, 2018, 16(2): 99–107.

[25] Wee J, Wei G W. Benchmarking AlphaFold3’s protein-protein complex accuracy and machine learning prediction reliability for binding free energy changes upon mutation [J]. ArXiv, 2024.

[26] Edwards M, Xu M, Joseph S. A simple procedure for bacterial expression and purification of the fragile X protein family [J]. Sci Rep, 2020, 10(1): 15858.

[27] Theler D, Dominguez C, Blatter M, et al. Solution structure of the Yth domain in complex with N6-methyladenosine Rna: a reader of methylated Rna [J]. Nucleic Acids Res, 2014, 42(22): 13911–9.

[28] Valverde R, Pozdnyakova I, Kajander T, et al. Fragile X mental retardation syndrome: structure of the KH1-KH2 domains of fragile X mental retardation protein [J]. Structure, 2007, 15(9): 1090–8.

[29] Mitrea D M, Kriwacki R W. Phase separation in biology; functional organization of a higher order [J]. Cell Commun Signal, 2016, 14: 1.

[30] Brangwynne C P, Eckmann C R, Courson D S, et al. Germline P granules are liquid droplets that localize by controlled dissolution/condensation [J]. Science, 2009, 324(5935): 1729–32.

[31] Su X, Ditlev J A, Hui E, et al. Phase separation of signaling molecules promotes T cell receptor signal transduction [J]. Science, 2016, 352(6285): 595–9.

[32] Antar L N, Afroz R, Dictenberg J B, et al. Metabotropic glutamate receptor activation regulates fragile x mental retardation protein and FMR1 MRNA localization differentially in dendrites and at synapses [J]. J Neurosci, 2004, 24(11): 2648–55.

[33] De Diego Otero Y, Severijnen L A, Van Cappellen G, et al. Transport of fragile X mental retardation protein via granules in neurites of PC12 cells [J]. Mol Cell Biol, 2002, 22(23): 8332–41.

[34] El Fatimy R, Davidovic L, Tremblay S, et al. Tracking the Fragile X Mental Retardation Protein in a Highly Ordered Neuronal RiboNucleoParticles Population: A Link between Stalled Polyribosomes and Rna Granules [J]. PLos Genet, 2016, 12(7): e1006192.

[35] Vernon R M, Chong P A, Tsang B, et al. Pi-Pi contacts are an overlooked protein feature relevant to phase separation [J]. Elife, 2018, 7.

[36] Tsang B, Arsenault J, Vernon R M, et al. Phosphoregulated Fmrp phase separation models activity-dependent translation through bidirectional control of MRNA granule formation [J]. Proc Natl Acad Sci U S A, 2019, 116(10): 4218–27.

[37] Myrick L K, Deng P Y, Hashimoto H, et al. Independent role for presynaptic Fmrp revealed by an FMR1 missense mutation associated with intellectual disability and seizures [J]. Proc Natl Acad Sci U S A, 2015, 112(4): 949–56.

[38] Collins S C, Bray S M, Suhl J A, et al. Identification of novel FMR1 variants by massively parallel sequencing in developmentally delayed males [J]. Am J Med Genet A, 2010, 152a(10): 2512–20.

[39] Zhang Q, Chen Z, Wang F, et al. Efficient DNA interrogation of SpCas9 governed by its electrostatic interaction with Dna beyond the Pam and protospacer [J]. Nucleic Acids Res, 2021, 49(21): 12433–44.

